# Transcription factors NFIA and NFIB induce cellular differentiation in high-grade astrocytoma

**DOI:** 10.1101/691303

**Authors:** Kok-Siong Chen, Jens Bunt, Caitlin R. Bridges, Zorana Lynton, Jonathan W.C. Lim, Brett W. Stringer, Revathi Rajagopal, Kum-Thong Wong, Dharmendra Ganesan, Hany Ariffin, Bryan W. Day, Linda J. Richards

## Abstract

Astrocytomas are composed of heterogeneous cell populations. Compared to grade IV glioblastoma, low-grade astrocytomas have more differentiated cells and are associated with a better prognosis. Therefore, inducing cellular differentiation may serve as a therapeutic strategy. The nuclear factor one (NFI) transcription factors are essential for normal astrocytic differentiation and therefore could be effectors of cellular differentiation in glioblastoma. We analysed expression of family members *NFIA* and *NFIB* using high-grade astrocytoma mRNA expression datasets, and with immunofluorescence co-staining. Their expression is reduced in glioblastomas and is associated with differentiated and mature astrocyte-like cells at a cellular level. Furthermore, induction of NFI expression is sufficient to promote cellular differentiation in patient-derived glioblastoma xenografts. Our findings indicate that NFI proteins may have an endogenous pro-differentiative function in astrocytomas, similar to their role in normal development. Overall, our study establishes a basis for further investigation of targeting NFI-mediated differentiation as a potential differentiation therapy.

## Introduction

Astrocytomas are characterised by high cellular heterogeneity, resulting in complex combinations of proliferating and differentiated cells (Aum et al., 2014; Louis et al., 2007; Patel et al., 2014). Compared with grade IV glioblastoma (GBM), lower grade astrocytomas consist predominantly of well-differentiated cells, and are associated with reduced aggressiveness and better prognosis (Louis et al., 2007). Hence, increasing cellular differentiation could render a tumour less aggressive and improve survival. This argument supports the notion that differentiation therapy could serve as a feasible therapeutic strategy. An example where this strategy has been remarkably successful is the treatment of acute promyelocytic leukaemia (APML) using retinoic acid (RA) (Huang et al., 1988). This success is the result of a multitude of studies investigating the role of RA in driving haematopoietic cell differentiation and targeting the fusion protein PML-RARα (Nowak et al., 2009).

Investigation of the response of GBM cell lines to inducers of differentiation did not yield promising results, as the responses appear transient and vary between cell lines (Campos et al., 2010; Caren et al., 2015; Choschzick et al., 2014). This is unsurprisingly given the endogenous function of these differentiation agents in normal brain cells. For example, RA has different functions depending on cell state, and is able to promote or inhibit differentiation at different stages of astrogliogenesis (Faigle et al., 2008). It is also ineffective in driving neuronal progenitor cell differentiation (Hadinger et al., 2009). One approach to overcoming such challenges is to target the endogenous mechanisms that are directly involved in producing mature, differentiated glial cells and re-deploy them in astrocytomas.

A promising candidate that may regulate differentiation in astrocytomas is the nuclear factor one (NFI) family of transcription factors. These proteins are essential for normal glial differentiation (Gobius et al., 2016; Namihira et al., 2009; Piper et al., 2010; Piper et al., 2009; Shu et al., 2003). During development, NFI are expressed in progenitor cells that give rise to various differentiated cell populations in the dorsal telencephalon (Bunt et al., 2017; Plachez et al., 2008). Their expression drives the differentiation of these cells (Gobius et al., 2016; Namihira et al., 2009; Piper et al., 2010), with NFIA and NFIB expression persisting in mature astrocytes (Chen et al., 2017a). In *Nfia* and *Nfib* knockout embryos, the generation of mature astrocytes is reduced, and progenitor cells remain in a proliferative state for a prolonged period (Piper et al., 2009; Shu et al., 2003). Overexpression of these genes is also sufficient to rapidly convert induced pluripotent stem cells into functional astrocytes *in vitro* (Caiazzo et al., 2015; Canals et al., 2018; Tchieu et al., 2019). These findings reflect the essential role of NFIA and NFIB as regulators of astrocytic differentiation.

In astrocytomas, higher expression levels of *NFI* correlate with lower grade tumours that are composed of more differentiated cells (Song et al., 2010; Stringer et al., 2016). Loss of heterozygosity (LOH) of *NFIB* as a consequence of chromosome 9p loss is a common occurrence in the tumour progression and is present in 40% of all GBM (Brat et al., 2004; Rasheed et al., 2002). The down-regulation of NFI expression may be a mechanism by which tumour cells evade differentiation and thereby remain proliferative. This is in line with studies which report that the *Nfi* loci are common insertion sites in insertional mutagenesis mouse models of gliomas, suggesting that reduced expression promotes tumorigenicity (Johansson et al., 2004; Vyazunova et al., 2014).

The induction of NFIA or NFIB in GBM cell lines *in vitro* induces the expression of astrocyte differentiation markers such as GFAP and FABP7 (Brun et al., 2009; Glasgow et al., 2014; Gobius et al., 2016; Stringer et al., 2016), indicating that overexpression of NFI might be sufficient to drive astrocytic differentiation. NFIA and NFIB function very similarly to drive glial differentiation in the developing brain (Gobius et al., 2016; Namihira et al., 2009; Piper et al., 2010). We demonstrated recently that NFIB overexpression *in vitro* reduced in GBM cell proliferation and inhibited growth when xenografted into mice (Stringer et al., 2016), although whether NFIA plays a similar role is unknown. Here, we investigate whether NFIA and NFIB drive astrocytic differentiation in astrocytomas *in vivo*. Using expression datasets and immunofluorescence co-staining, we demonstrate that the endogenous expression of NFIA and NFIB positively correlate with each other, and that their expression in astrocytomas is associated with the expression of differentiated cell markers. Furthermore, by manipulating their expression in patient-derived GBM xenografts *in vivo*, we reveal that induced expression of either NFI is sufficient to promote tumour cell differentiation. These findings suggest that the NFI-pathway is a promising therapeutic target to induce differentiation in astrocytomas.

## Results

### *NFIA* and *NFIB* expression is decreased in grade IV GBM

To compare the expression of *NFIA* and *NFIB*, and their relationship in different grades of astrocytomas, we analysed seven large mRNA expression datasets of human astrocytoma samples. Both *NFIA* and *NFIB* expression were reduced in GBM as compared to grade I-III astrocytomas (Fig. 1a), demonstrating that the expression of both *NFIA* and *NFIB* is higher in tumours with more differentiated tumours, and lower in poorly differentiated tumours. We also observed a positive correlation between *NFIA* and *NFIB* expression in GBM samples across ten datasets (Fig. 1b; Supplementary Table 2). Although the strength of the correlation coefficient varied among datasets, this variability may be explained by LOH of *NFIB* observed in a subset of GBM samples (Brat et al., 2004; Rasheed et al., 2002). Separating tumour samples based on *NFIB* copy number significantly improved the correlation coefficient between *NFIA* and *NFIB* expression in samples where these data were available (Fig. 1c). The trendlines representing the *NFIB*-diploid and haploinsufficient tumour samples had a similar correlation coefficient and were separated by approximately one ^2^log unit. This meant that when comparing between samples with similar *NFIA* expression, *NFIB* expression was halved in *NFIB*-haploinsufficient samples, as would be expected when one *NFIB* allele is lost. Similar analyses for *NFIA* haploinsufficiency, which is rarely observed in GBM, resulted in only a small improvement in the correlation coefficient (Supplementary Table 3).

**Figure 1.**
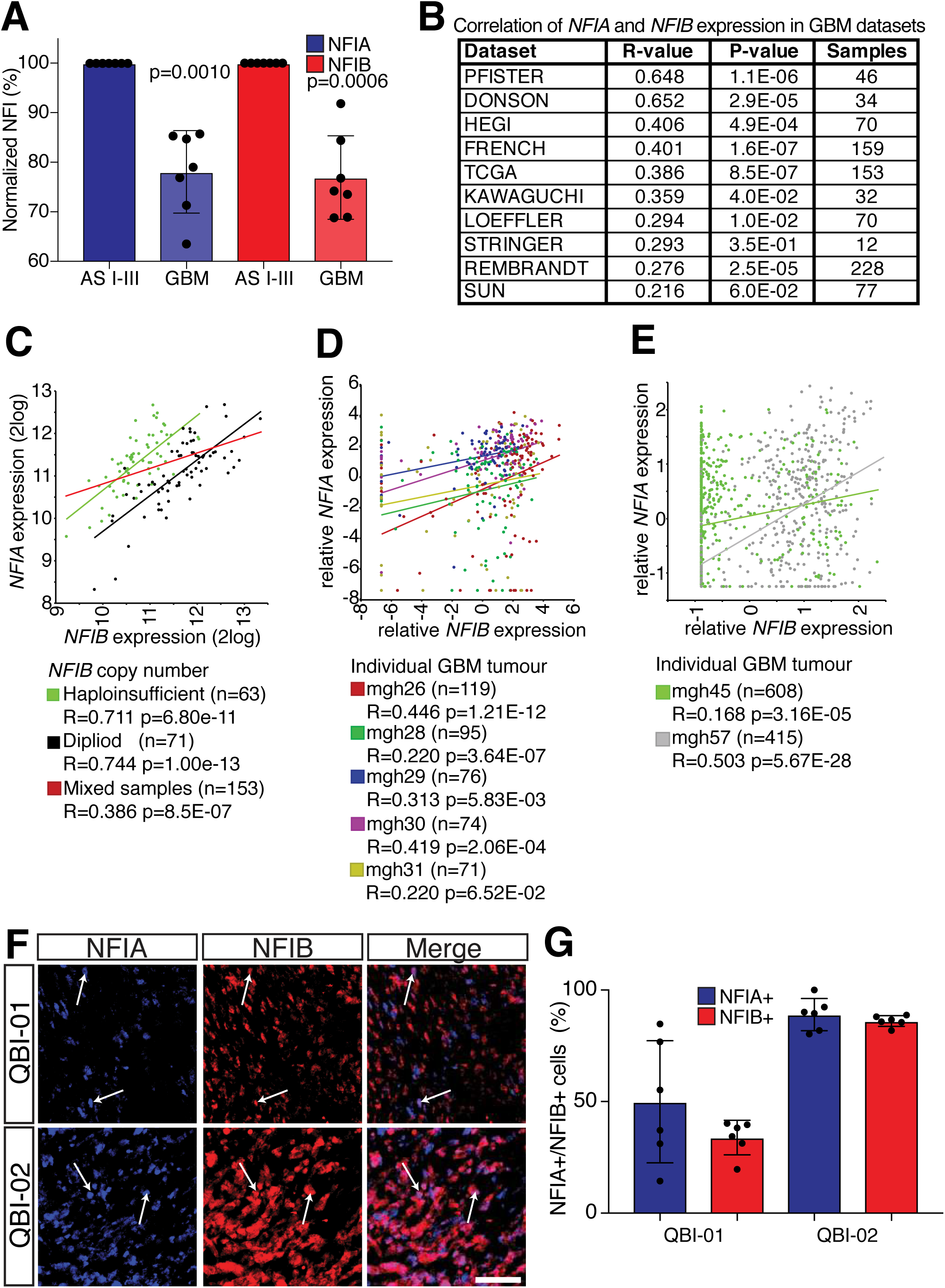
*NFIA* and *NFIB* share a similar expression pattern in astrocytoma. **a** Based on seven mRNA expression datasets, the expression of both *NFIA* and *NFIB* decreases in grade IV GBM as compared to grade I-III astrocytoma (Kruskal-Wallis test). **b** *NFIA* and *NFIB* expression positively correlate with each other within all analysed mRNA expression datasets of GBM. **c** By separating GBM samples in the TCGA mRNA dataset based on *NFIB* copy number, the correlation between *NFIA* and *NFIB* expression improves, resulting in higher correlation coefficients for both groups. **d** and **e** *NFIA* and *NFIB* expression positively correlate in single-cell expression data of seven GBM tumours. **f** Immunofluorescence co-staining of NFIA and NFIB proteins (arrows) in two patient-derived GBM xenograft tissues. **g** Quantification of NFIA+/NFIB+ cells as a percentage of all NFIA+ or NFIB+ cells. Scale bar in **f** = 50 *µ*m. Error bars represent the standard deviation of the mean.

### *NFIA* and *NFIB* are co-expressed at the cellular level in GBM tumours

The correlation between *NFIA* and *NFIB* expression in tumour samples does not necessarily mean that these genes are co-expressed within the same cells. We investigated this by first examining *NFIA* and *NFIB* expression in GBM tumours analysed by single cell RNA-seq (Patel et al., 2014; Venteicher et al., 2017).

Similar to our observations with whole tumour samples, we observed that the expression of *NFIA* and *NFIB* positively correlated with each other at the single-cell level (Fig. 1d and e; Supplementary Table 4). To validate this at the protein level, we then performed immunofluorescence co-staining of NFIA and NFIB in patient-derived GBM xenografts (Supplementary Table 1a and 5). Quantification was performed with two biological replicates in two non-necrotic 500 × 500 *µ*m ROIs per tumour. In the QBI-01 xenograft, which represents a tumour with fewer NFI-positive cells in general, half of all NFIA-positive cells co-stained for NFIB, whereas over a third of the NFIB-positive cells were also NFIA-positive (Fig. 1f and g). The co-expression of NFIA and NFIB was observed in more cells for the QBI-02 xenograft, in which at least 80% of cells expressed both NFIA and NFIB. We therefore conclude that NFIA and NFIB show a high degree of co-expression within the same cells in GBM tumours.

### *NFIA* and *NFIB* expression correlates with astrocytic differentiation genes in GBM cells

GBM tumours consist of a heterogenous mix of cell types (Aum et al., 2014; Patel et al., 2014). Highly aggressive tumours that consist predominantly of proliferating cells still harbour differentiated tumour cells that are not proliferating (Louis et al., 2007). Given their role as regulators of cell differentiation during development, we hypothesized that *NFIA* and *NFIB* are associated with genes that correspond with a differentiated cell state. To investigate this, we performed gene ontology (GO) analyses on gene sets that were independently derived from each of the three largest GBM mRNA expression datasets (Supplementary Table 6 and 7). We generated four gene sets for each mRNA expression dataset, with each set consisting of 500 genes. The first two sets consisted of the top genes that positively correlated with *NFIA* or *NFIB*, respectively, while the remaining two sets consisted of the top genes that showed negative correlation with *NFIA* or *NFIB*. Each set was then subjected to GO analyses to identify which GO Biological Process terms were enriched.

The six gene sets that were positively correlated with *NFIA* or *NFIB* (two sets for each of the mRNA expression datasets) returned a combined total of 345 unique GO terms (p<0.001) (Fig. 2a; Supplementary Table 7). 114 of these terms (33%) occurred in at least four of the six positively correlated gene sets, but were completely absent from the six gene sets that are negatively correlated with *NFIA* or *NFIB*. These included GO terms associated with transcription, metabolic processes and development. 14 of these terms were shared between all six positively correlated gene sets. These are likely to represent genes that are co-expressed in NFIA+ and NFIB+ cells, and are enriched for terms representative of neurodevelopment and cell differentiation processes (Fig. 2b). Overall, these findings are in line with the strong co-expression of NFIA and NFIB observed in these datasets (Fig. 1b-g), and indicative of their overlapping function in regulating differentiation. Notably, no GO terms associated with proliferation were enriched in any of the positively or negatively correlated gene sets, suggesting that *NFI* expression is indicative of tumour differentiation state and not proliferative potential.

**Figure 2.**
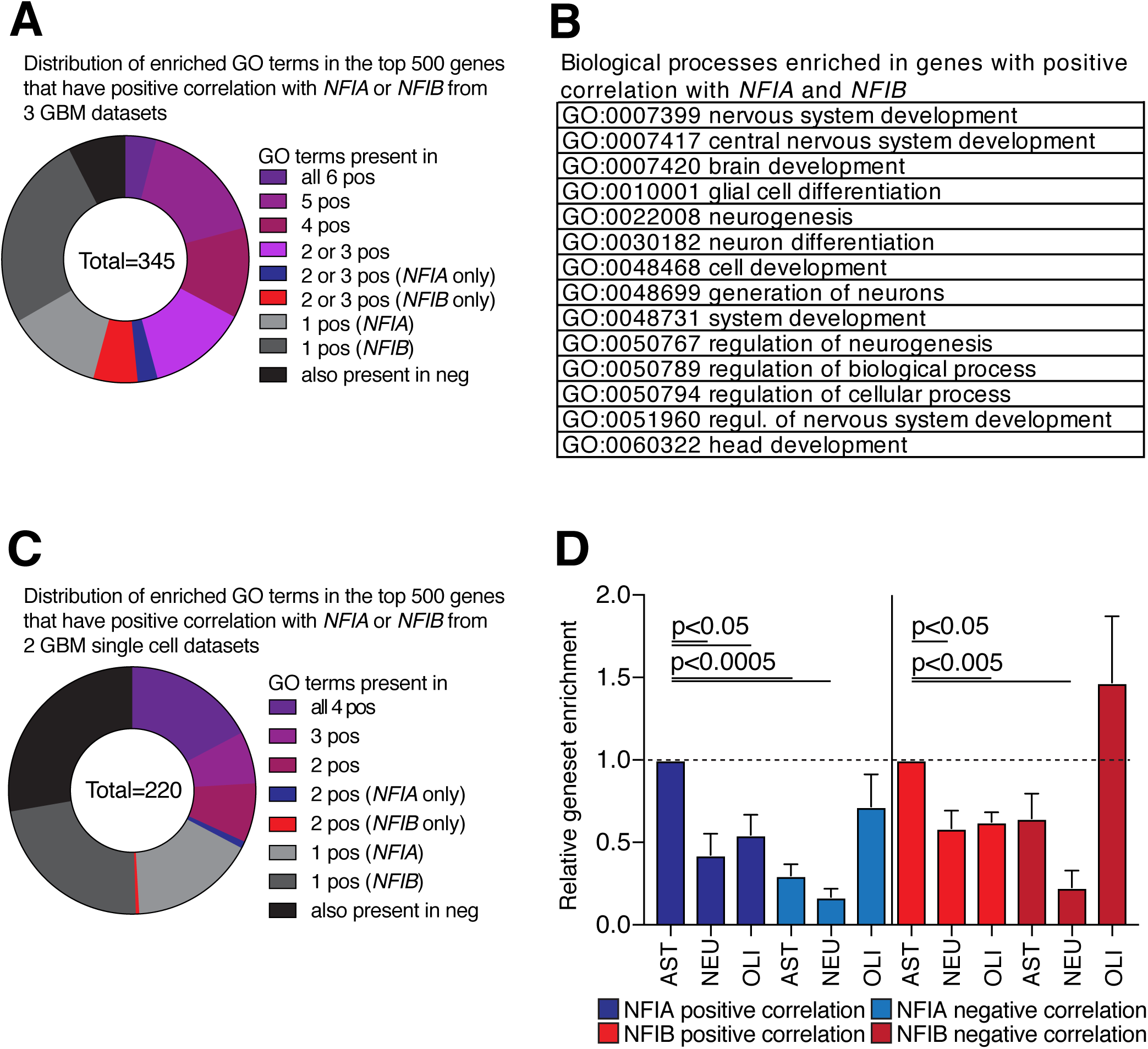
*NFI* expression in GBM is associated with neurodevelopment and mature astrocytic genes. **a** Gene ontology (GO) analyses were performed on gene sets generated from three large GBM expression datasets. Four gene sets were generated for each large dataset, representing the top 500 genes that positively correlate or negatively correlate with *NFIA* or *NFIB*. Of the 345 GO Biological Process terms obtained from the six positive gene sets (pos), 45% were shared by both *NFIA*- or *NFIB*-correlated gene sets, but were absent from all of the negatively correlated gene sets (neg). **b** The 14 GO terms present in all six gene sets that positively correlated with *NFI* expression. **c** Similar GO analyses were performed on two large GBM single-cell mRNA-seq datasets. Of the 220 GO Biological Process terms obtained from the four gene sets representing the *NFIA* or *NFIB* positively correlated genes, 32% of the GO terms were shared between *NFIA*- and *NFIB*-correlated genes in both datasets. **d** Relative enrichment of genes associated with mature astrocytic (AST), oligodendrocytic (OLI) and neuronal (NEU) signatures derived from the positive and negative gene sets that were generated for all eight datasets. *NFI* positively correlated genes were associated with mature astrocytic genes. Statistical significance was determined using a paired one-way ANOVA with Bonferroni’s multiple comparison correction.

We next performed similar analyses using the two largest single-cell mRNA-seq datasets derived from individual GBM tumours. Other single-cell datasets were excluded, as these did not contain sufficient cells for reliable correlation analyses. The four gene sets representing *NFI* positively correlated genes were enriched for a combined total of 220 GO terms (p < 0.00005). 70 of these terms (32%) were shared at least once between the *NFIA* and *NFIB* gene sets, but were completely absent from the gene sets representing negatively correlated genes (Fig. 2c; Supplementary Table 8 and 9). Similar to our analyses of whole GBM tumours, the 38 GO terms shared between all four positively correlated gene sets represented neurodevelopmental processes, such as neurogenesis, gliogenesis, and cell differentiation (Supplementary Table 9). These findings also held true for gene expression datasets of two GBM cell lines collected from different passages and culture conditions (Supplementary Table 10 and 11) (Lee et al., 2006), further supporting the premise that *NFI* expression is a marker of differentiated cells.

To determine whether the correlated gene sets were associated with specific cell types, we compared these sets with gene expression signatures that represent mature astrocyte, oligodendrocyte or neuron-specific genes as reported by three independent groups (Supplementary Table 12) (Cahoy et al., 2008; Lein et al., 2007; McKenzie et al., 2018). The *NFIA* and *NFIB* positively correlated gene sets were associated with astrocytic genes (Fig. 2d). In contrast, genes that negatively correlated with *NFIA* or *NFIB* were enriched for oligodendrocytic markers. This suggests that *NFI-*expressing tumour cells are likely to resemble mature astrocytes rather than oligodendrocytes or neurons.

### NFIA and NFIB proteins mark non-proliferating, differentiated tumour cells

To determine whether NFI expression is associated with astrocytic markers in GBM tumours *in vivo*, we performed immunofluorescence co-staining on sections derived from five primary GBM samples, five xenografts derived from GBM cell lines and three patient-derived xenograft tumours (Fig. 3; Supplementary Fig. 1 and 2) (Stringer et al., 2019). All samples expressed both NFIA and NFIB, and displayed co-staining of markers of astrocytic differentiation. Cell counts demonstrated that the majority of NFIA- or NFIB-positive cells (∼80%) were also positive for GFAP or S100B (Fig. 4; Supplementary Table 13 and 14). In contrast, only one third of the NFIA- or NFIB-positive cells co-stained with the proliferation marker Ki67. Hence, NFI expression is associated with the expression of mature astrocytic markers.

**Figure 3.**
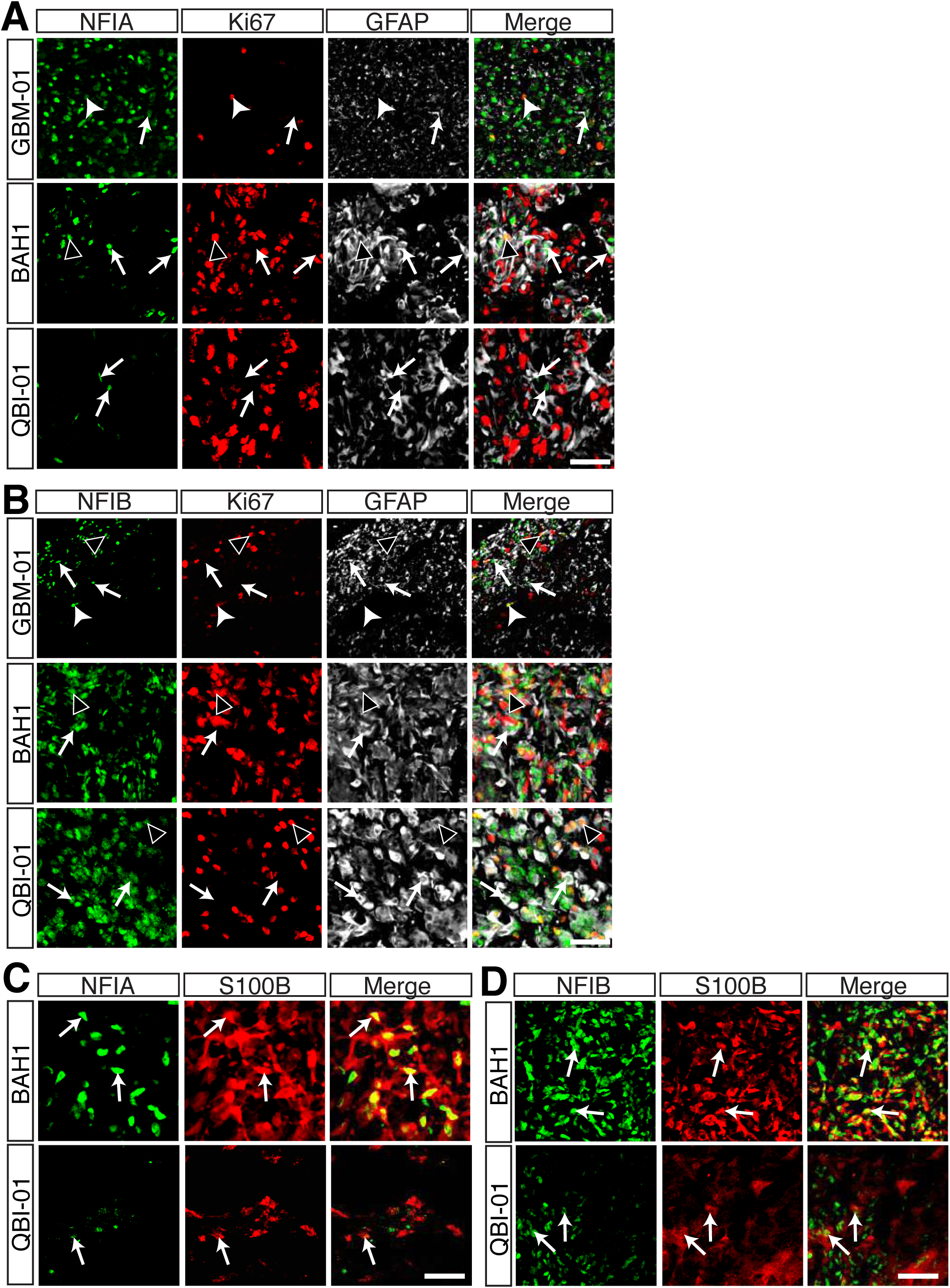
NFI-expressing cells in GBM are predominantly associated with the astrocytic differentiation marker GFAP. **a** and **b** Representative images of co-staining of NFIA (**a**) or NFIB (**b**) with GFAP and the proliferation marker Ki67. **c** and **d** Co-staining of NFIA (**c**) or NFIB (**d**) with the astrocytic marker S100B in GBM tissues. Closed arrowhead: NFI, GFAP and Ki67 co-localisation; open arrowhead: NFI and Ki67 co-localisation; arrow: NFI and GFAP co-localization. Scale bar (**a, b, c, d**) = 50 *µ*m.

**Figure 4.**
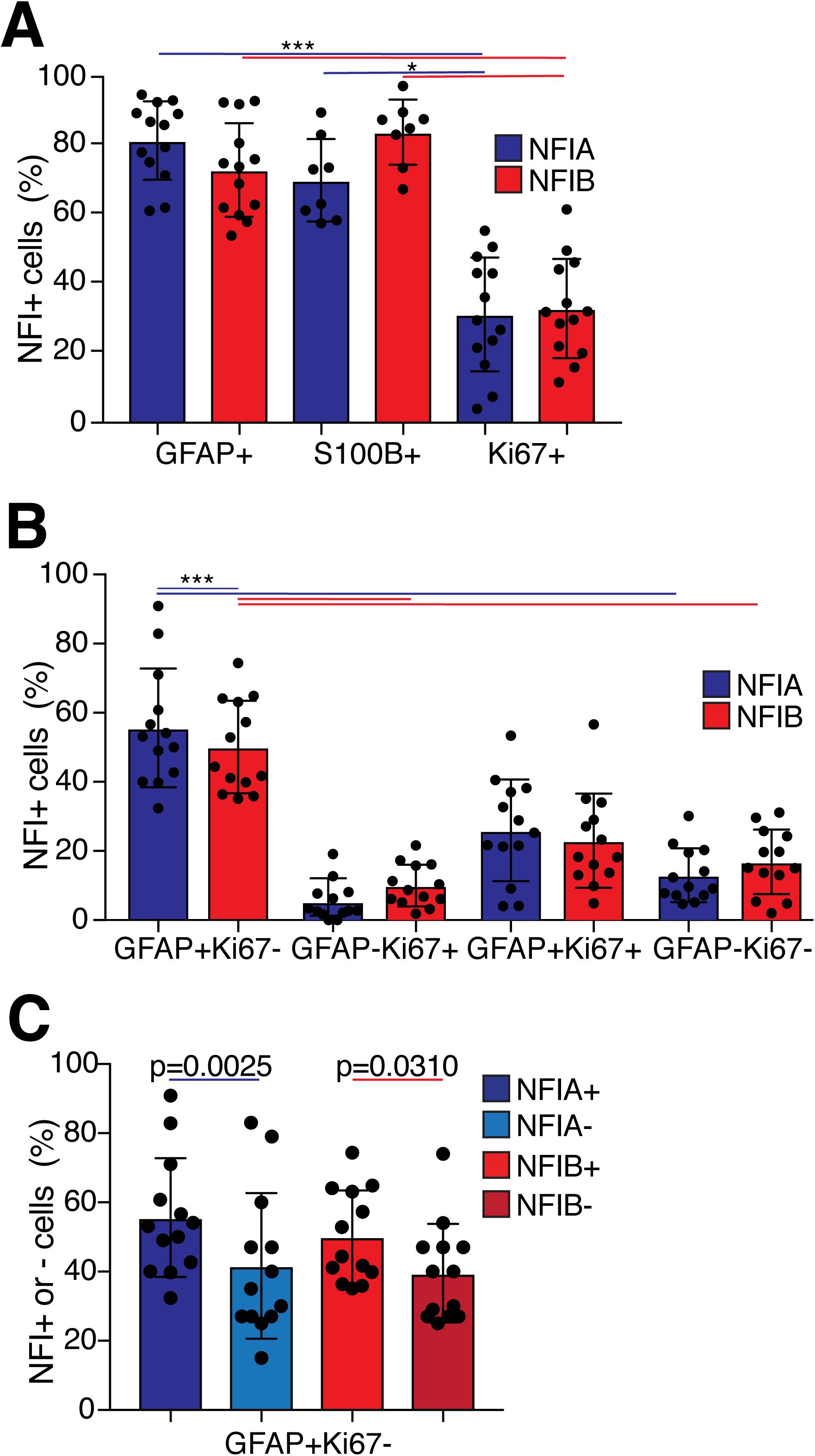
NFI-expressing cells in GBM are predominantly associated with astrocytic differentiation genes. **a** Quantification of NFIA or NFIB co-staining with astrocytic and proliferation markers, demonstrating the percentage of NFIA+ and NFIB+ cells co-stained for GFAP, S100B or Ki67. **b** The percentage of NFIA+ and NFIB+ cells co-stained with GFAP and Ki67, respectively. **c** GFAP+/Ki67-cells as a percentage of NFI positive or negative cells. Error bars represent the standard deviation of the mean. Statistical significance was determined using a paired one-way ANOVA with Bonferroni’s multiple comparison correction; \**p* <0.05, *** *p* < 0.0005.

To further delineate whether NFI-expressing cells are fully differentiated, we analysed the combined co-staining of GFAP and Ki67 (Fig. 3 and 4; Supplementary Fig. 1 and 2; Supplementary Table 13 and 14). No difference in the co-staining pattern was observed between NFIA-positive or NFIB-positive cells. Over half of the NFI-positive cells co-expressed only GFAP (Fig. 4c), indicative of fully differentiated cells, whereas less than a third of these cells were both GFAP- and Ki67-positive. The identity of the latter cells is questionable, but could perhaps represent cells that have just begun differentiating. We also observed that a subset of GFAP-positive cells was devoid of NFIA or NFIB expression. However, we were unable to determine whether both NFIA and NFIB were concurrently absent in these cells, as our NFIA and NFIB antibodies were derived from the same host species.

### Expression of NFIA or NFIB induces tumour cell differentiation

NFI expression is indicative of differentiated cells that express mature astrocytic markers in GBM, but whether these transcription factors can actually drive the differentiation of proliferating cancer cells remains unclear. Overexpression of either NFIA or NFIB is sufficient to induce the expression of astrocytic markers *in vitro* (Brun et al., 2009; Canals et al., 2018; Stringer et al., 2016), but this remains to be demonstrated for tumours *in vivo*. In addition, no study has yet to compare both NFIA and NFIB under similar conditions. To investigate whether NFI expression can drive astrocytic differentiation *in vivo*, we introduced NFIA or NFIB overexpression plasmids via *in vivo* electroporation into subcutaneous xenografts. As a proof of concept, we electroporated xenografts of the U251 GBM cell line, which is responsive to NFIB induction *in vitro* (Brun et al., 2009). Electroporated xenografts were sectioned and co-labelled for GFAP, Ki67 and green fluorescent protein (GFP), which is indicative of cells that were successfully electroporated (Fig. 5a). The overall number of GFAP-positive cells increased in NFIB-electroporated xenografts as compared to xenografts electroporated with the control plasmid (Fig. 5b; Supplementary Table 15). Specifically, we observed a concurrent decrease in GFAP-negative, Ki67-positive cells and a subsequent increase in GFAP-positive, Ki67-negative cells, indicating that proliferative cells transitioned into a differentiated state upon NFIB overexpression. A small but insignificant increase in the number of GFAP-positive, Ki67-positive cells was observed, suggesting that these could represent proliferative cells that are transitioning to a differentiated state.

**Figure 5.**
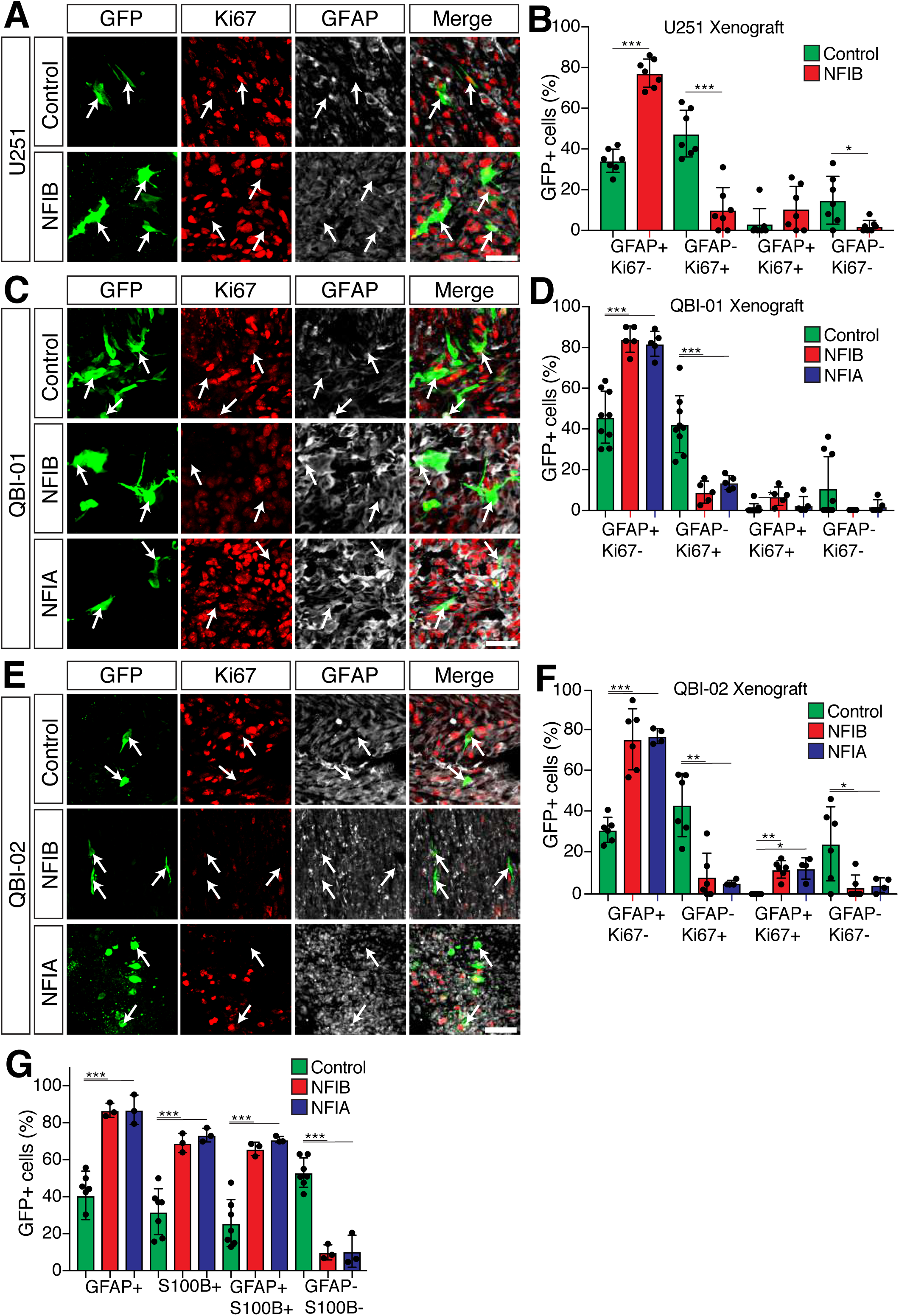
*In vivo* electroporation of GBM xenografts with NFI overexpression constructs. **a, c** and **e** Examples of GFP+ cells (arrows) as a marker of cells electroporated with control, NFIA or NFIB overexpression plasmids in U251 (**a**), QBI-01 (**c**) and QBI-02 (**e**) xenografts. Sections were also stained for the proliferation marker Ki67 (red) and the astrocytic differentiation marker GFAP. **b, d** and **f** Quantification of co-localization of the xenografts represented as a percentage of the GFP+ cells in U251 (**b**), QBI-01 (**d**) and QBI-02 (**f**) xenografts electroporated with control, NFIA or NFIB overexpression plasmids. **g** Quantification of co-localization of GFP+ cells in QBI-02 with S100B and GFAP. Error bars represent the standard deviation of the mean. Statistical significance was determined using a Welch’s t-test; \**p* <0.05, \*\**p* <0.005, *** *p* < 0.0005. Scale bars (**a, c, e**) = 50 *µ*m.

To determine whether overexpression of NFIA or NFIB exerts the same effect, we electroporated NFIA, NFIB or control plasmids into xenografts representing two patient-derived xenograft lines (Fig. 5c and e; Supplementary Table 15). These lines responded similarly to either NFIA or NFIB overexpression, with the number of GFAP-positive, Ki67-negative cells increasing to approximately 80% in NFIA- or NFIB-electroporated xenografts (Fig. 5d and f). We observed a concurrent reduction in the number of cells that were GFAP-negative, Ki67-positive, and GFAP-negative, Ki67-negative, while GFAP-positive, Ki67-positive cells increased. Immuno-labelling of the QBI-02 xenografts with S100B, another marker expressed in mature astrocytes, returned similar findings (Fig. 5g; Supplementary Fig. 4a and b). Thus, we conclude that either NFIA or NFIB is sufficient to induce astrocytic differentiation in GBM.

## Discussion

In this study, we have demonstrated that both NFIA and NFIB function as regulators of astrocytic differentiation in astrocytoma. NFI expression is higher in lower grade astrocytomas, and the expression of NFIA and NFIB in GBM is typically associated with genes representing the mature astrocytic state. Furthermore, both these transcription factors are co-expressed in GBM cells that also express mature astrocytic markers. Most importantly, *in vivo* overexpression of either NFIA or NFIB in xenografts is sufficient to drive proliferative cells towards astrocytic differentiation. Hence, NFIA and NFIB retain their developmental role as inducers of astrocytic differentiation in astrocytoma, and could therefore act as tumour suppressors within a therapeutic context.

Although strong evidence exists to suggest that NFIB acts as a tumour suppressor in astrocytoma (Johansson et al., 2004; Patel et al., 2014; Stringer et al., 2016; Vyazunova et al., 2014), the role of NFIA remains disputable (Glasgow et al., 2013; Johansson et al., 2004; Lee et al., 2014; Patel et al., 2014; Song et al., 2010; Vyazunova et al., 2014). *In vitro* experiments suggest that NFIA promotes rather than suppresses cancer cell proliferation (Lee et al., 2014). However, our findings demonstrate that both NFIA and NFIB function as tumour suppressors as they drive proliferating cells to differentiate towards an astrocytic fate *in vivo*. Whether they share similar functions in other tumours requires further study, but this is unlikely to be the case. These transcription factors are widely expressed in many tissues during development, and both oncogenic and tumour suppressive roles have been reported which may be tissue-specific (Chen et al., 2017b). For instance, NFIA plays a vital role in the development of oligodendrocytes (Kang et al., 2012; Wong et al., 2007), with the expression of NFIA, but not NFIB, being retained in adult oligodendrocytes (Chen et al., 2017a). In line with this, oligodendrogliomas tend to demonstrate lower *NFIA* expression than astrocytomas, in contrast to *NFIB* (Song et al., 2010). This lower expression may be associated with the partial loss of chromosome 1p31, which is more often observed in oligodendrogliomas than in astrocytomas (Houillier et al., 2010). Interestingly, overexpression of NFIA in a mouse model of oligodendroglioma resulted in tumours resembling astrocytomas (Glasgow et al., 2014). In spite of this, as this study did not investigate whether these cells remained proliferative, another possible interpretation of the findings is that NFIA overexpression caused the cells to differentiate towards the astrocytic lineage, similar to our observations.

NFI proteins have also been implicated as tumour suppressors in other brain cancers, such as the SHH-subtype of medulloblastomas (Genovesi et al., 2013; Wu et al., 2012). This role was strongly corroborated using a mouse model of SHH-subtype medulloblastoma with heterozygous deletion of *Nfia* (Genovesi et al., 2013). Compared to normal *Nfia* expression, loss of one allele increased tumour incidence and decreased tumour latency. Hence, NFI may broadly function as inducers of differentiation in different brain tumour cell types.

The identification of the *Nfi* loci as common insertion sites in insertional mutagenesis mouse models demonstrates that reduced *Nfi* expression contributes to glioma tumorigenesis (Johansson et al., 2004; Vyazunova et al., 2014). However, how tumour cells suppress *NFI* expression in astrocytomas *in vivo* to evade differentiation remains unclear. As mutations of the *NFI* genes are rare, haploinsufficiency of *NFIA* or *NFIB* appears to be the most common pathway through which their expression is reduced. It is also likely that *NFI* expression is regulated on a transcriptional level in proliferating tumour cells that have evaded differentiation commitment. Unfortunately, little is known about the upstream regulators of *NFI* during normal astrogliogenesis or in astrocytoma, so this requires further investigation. However, due to the strong positive correlation between *NFIA* and *NFIB* expression, it is likely that both have similar upstream regulators. A recent study proposed that *NFIA* expression could be mediated by TGFβ in normal gliogenesis (Tchieu et al., 2019). Whether reduced *NFIA* expression in GBM is due to ectopic TGFβ signalling remains to be elucidated. Aside from this, post-transcriptional regulation may also contribute to *NFI* down-regulation. For example, the expression of the microRNAs miR-124 and miR-129 positively correlates with increased glioma grade, but inversely correlates with *NFIB* expression (Ho et al., 2013; Silber et al., 2008). Indeed *in vitro* experiments have demonstrated that microRNAs regulate *NFI* expression in astrocytoma cell lines, but their significance *in vivo* remains to be determined. Greater emphasis on understanding how *NFI* is suppressed in tumour cells is required for effective therapeutic manipulation of the NFI-mediated differentiation pathway.

In addition to direct manipulation of the NFI pathway as a potential differentiation therapy, increased expression of NFI may also act as a biomarker indicative of differentiation when testing novel therapeutic agents or for diagnostic purposes. It is not known whether NFI expression was induced and sustained for the differentiation agents tested (Campos et al., 2010; Caren et al., 2015; Choschzick et al., 2014). The abundance of NFI proteins could even indicate whether a tumour would be more prone to induction of differentiation for differentiation therapies instead of, or in combination with, the current standard treatment regime. Nevertheless, any diagnostic or prognostic value of NFIA or NFIB to clinical management will require further investigation.

In conclusion, our study demonstrates that both NFIA and NFIB play a direct role in inducing tumour differentiation in astrocytoma. Given that tumours with fewer differentiated cells are associated with a poorer clinical outcome, a deeper understanding of the NFI-mediated differentiation mechanisms may reveal a potential therapeutic strategy to reduce the proliferative potential of GBM cells via differentiation, and thereby improve patient survival.

## Material and Methods

### *In silico* analyses

To assess gene expression in human glioma samples, public expression datasets GSE50161 (Griesinger et al., 2013), GSE36245 (Sturm et al., 2012), GSE16011 (Gravendeel et al., 2009), GSE7696 (Murat et al., 2008), GSE43378 (Kawaguchi et al., 2013), GSE53733 (Reifenberger et al., 2014), GSE57872 (Patel et al., 2014), GSE108474 (Gusev et al., 2018), GSE118793, GSE4290 (Sun et al., 2006), GSE89567 (Venteicher et al., 2017), and TCGA (Brennan et al., 2013) were analysed and visualised using the R2: microarray analysis and visualization platform (http://r2.amc.nl) as previously described (Bunt et al., 2010). Gene set enrichment analyses were performed using The Database for Annotation, Visualization and Integrated Discovery (DAVID) 6.8 online tool (Huang da et al., 2009a, 2009b).

### Animals

All breeding and experiments were performed at The University of Queensland in accordance with the Australian Code of Practice for the Care and Use of Animals for Scientific Purposes, and with the approval of the University of Queensland Animal Ethics Committee. Animals were housed on a 12 hour dark/light cycle with water and food provided *ad libitum*. The NOD.CB17-Prkdc^scid^/Arc (NOD-SCID) mice used for patient-derived GBM xenograft experiments were obtained from the Australian Animal Resources Centre.

### Generation and collection of GBM xenografts

The establishment of xenografts was performed as previously described with minor modifications (Carlson et al., 2011). For patient-derived GBM xenografts (QBI-01, QBI-02, and QBI-03), fresh GBM patient tumour samples were obtained from the Wesley Medical Research BioBank with approval from the University of Queensland Human Ethics Committee. Upon receipt, samples were disaggregated using a scalpel blade and mixed with extracellular matrix (ECM) gel (Sigma-Aldrich, St. Louis, Missouri) in a 1:1 ratio on ice. Cell line-derived xenografts were generated from U251 (Ponten et al., 1968) and patient-derived GBM cell lines BAH1, SJH1, RKI1, and WK1 obtained from the QIMR Berghofer Medical Research Institute (Day et al., 2013; Stringer et al., 2019). The U251 GBM cell line was maintained *in vitro* in Dulbecco’s Modified Eagle Medium supplemented with 10% (v/v) foetal bovine serum (FBS) (Life Technologies, Carlsbad, California), whereas patient-derived cell lines were cultured as adherent monolayers in ECM gel-coated vessels using RHB-A stem cell culture medium (Life Technologies) supplemented with 20 ng/mL epidermal growth factor (Life Technologies), 10 ng/mL fibroblast growth factor 2 (Life Technologies), and StemPRO NSC SFM (Life Technologies) as previously described (Pollard et al., 2009). All cell lines were incubated at 37°C with 5% CO_2_. To dissociate adherent cells, 1 mL of 0.25% pre-warmed trypsin (Life Technologies) was added to the T25 culture flask (Corning) for U251, whereas 1 mL pre-warmed StemPro^®^ Accutase® (Life Technologies) was used for patient-derived cell lines. The dissociation reagent was then inhibited following cell dissociation using Trypsin Inhibitor (Life Technologies). The cell suspension was centrifuged at 200 relative centrifugal force (RCF) for 5 min and the cell pellet was resuspended with ECM gel in a 1:1 ratio on ice. 50-200 *µ*L of the tumour cells / ECM gel mixture was injected using a 16-21 G needle into the flank of NOD/SCID mice anaesthetised with methoxyflurane (Medical Developments International, Scoresby, Australia).

### *In vivo* electroporation of GBM xenografts

NFI expression in established GBM xenografts was driven using the piggyBac system as described previously (Chen et al., 2012). pPBCAGIG, pPBCAGIG-NFIB or pPBCAGIG-NFIA donor plasmids were co-electroporated with pCAG-PBase as a helper plasmid into established xenografts of 0.1-0.2 cm^3^ (Hsieh et al., 2003; Tsurushima et al., 2008). pPBCAGIG was generated by cloning the CAG-ires-eGFP cassette from pCAGIG into pPBCAG-eGFP. Subsequently, HA-tagged *Nfia* or HA-tagged *Nfib* (Piper et al., 2014; Stringer et al., 2016) was cloned into this vector to generate pPBCAGIG-NFIA or pPBCAGIG-NFIB. The pCAG-PBase and pPBCAG-eGFP plasmids were kind gifts from Joseph LoTurco, University of Connecticut. Electroporations were performed on xenografted mice anaesthetised with methoxyflurane and placed on a heat pad. 20 *µ*l of the mixture of 0.5 *µ*g/*µ*l helper plasmid and 0.5 *µ*g/*µ*l donor plasmid in distilled water was injected into the centre of the xenograft with a 31 G needle. Electroporation was performed using 3mm electrodes (Nepagene, Ichikawa, Japan), placed on either side of the xenograft. Six pulses, each of 150V for a duration of 50ms, were administered with a frequency of 1 pulse/s using the ECM 830 electroporator (BTX Harvard Apparatus, Holliston, Massachusetts). Another 6 pulses were delivered in the opposite direction by switching the positions of the electrodes. Animals were sacrificed by cervical dislocation four days after electroporation and resected xenografts were fixed in 4% (w/v) paraformaldehyde (PFA) in phosphate buffered saline (PBS) (Lonza, Basel, Switzerland) at 4°C for approximately one week. Prior to sectioning, the PFA solution was replaced with 5% formalin diluted in absolute ethanol, and the xenograft was incubated at room temperature for a further 6 h.

### Tissue processing

Following post-fixation, xenografts and mouse brain tissue were embedded in 3-4% (w/v) Difco Noble agar (BD Biosciences, San Jose, California) in distilled water, and sectioned at 50 *µ*m on a vibratome (Leica, Wetzlar, Germany). Sections were stored in PBS supplemented with 0.2% (w/v) sodium azide at 4°C or mounted directly onto SuperfrostPlus slides (Menzel-Gläser, Braunscheig, Germany), and dried at room temperature until fully adhered. Adherent sections were post-fixed with 4% PFA for 10 min and washed with PBS for 5 min before further processing.

Prior to antigen retrieval, mounted xenograft sections and paraffin-embedded human GBM tissue sections (obtained from the University of Malaya Biobank with approval from the University of Queensland Human Ethics Committee) were dehydrated in a series of ethanol (70% and 100%) washes, defatted in xylene (VWR, Radnor, Pennsylvania) and rehydrated through a series of ethanol (100% and 70%) washes. Antigen retrieval was performed using an antigen decloaking chamber (Biocare Medical, Pacheco, California). Mounted sections were heated to 125°C for 4 minutes at 15 psi in 10 mM sodium citrate buffer (pH 6.0) with 0.05% (v/v) Tween 20. Following antigen retrieval, sections were washed with PBS prior to immunohistochemistry. All tissues used are listed in Supplementary Table 1a.

### Immunohistochemistry

Fluorescence immunohistochemistry was performed as previously described with minor modifications (Plachez et al., 2008). After antigen retrieval, sections were incubated for 2 h in blocking solution containing 10% (v/v) normal donkey or goat serum (Jackson Laboratories, Bar Harbor, Maine), 0.9% (v/v) hydrogen peroxide (Chem-Supply, Gillman, Australia) and 0.2% (v/v) Triton X-100 (Sigma-Aldrich) in PBS. Sections were then incubated overnight with primary antibodies diluted in 2% (v/v) serum and 0.2% (v/v) Triton X-100 in PBS. The primary antibodies used are listed in Supplementary Table 1b. Sections were washed in PBS for 3×20 minutes before incubating with secondary antibody.

For standard fluorescence immuno-labelling, the secondary antibodies used were either donkey anti-rabbit IgG, donkey anti-mouse IgG, donkey anti-goat IgG or donkey anti-chicken IgG Alexa Fluor 488, 555, or 633 (1:500; Invitrogen, Carlsbad, California) diluted in PBS containing 0.2% v/v Triton X-100, and incubated for 3 h in a light-protected humidified chamber. For fluorescence amplification, sections were incubated with biotinylated donkey anti-rabbit IgG, anti-chicken IgG or anti-mouse IgG secondary antibody (1:500; Jackson Laboratories) for 1 hour. After 3×10 min washes with PBS, the sections were then incubated with Strepavidin-Alexa Fluor 647 conjugate (1:500; Invitrogen) diluted in 0.2% (v/v) Triton X-100 in PBS for 1 h.

Following immune-labelling, sections were washed for 3×10 min with PBS and incubated in 4’,6-diamidino-2-phenylindole, dihydrochloride (DAPI; 1:1000; Invitrogen) in 0.2% (v/v) Triton X-100 in PBS for 5 min. Sections were then washed for 3×10 min with PBS and cover-slipped using ProLong Gold anti-fade reagent with (Invitrogen) as a mount solution.

### Image acquisition and analysis

High resolution fluorescence images were acquired using a Marinas spinning-disk confocal system (3I, London, UK) consisting of an Axio Observer Z1 (Carl Zeiss) equipped with a CSU-W1 spinning-disk head (Yokogawa Corporation of America, Sugar Land, Texas), ORCA-Flash 4.0 v2 sCMOS camera (Hamamatsu Photonics, Japan), 20x 0.8 NA PlanApo objective, captured with SlideBook 6.0 (3I, Denver, Colorado), or a Diskovery spinning disk confocal system (Spectral Applied Research, Ontaria, Canada) built around a Nikon TiE body and equipped with two sCMOS cameras (Andor Zyla 4.2, 2048 × 2048 pixels) and captured with Nikon NIS software (Nikon, Tokyo, Japan). Images were pseudo-coloured to permit overlay, cropped, sized, and contrast-brightness enhanced for presentation with Photoshop and Illustrator software (Adobe, San Jose, California).

Semi-automated cell counting of fluorescence images was performed using Imaris software (Bitplane, Belfast, United Kingdom), version 8.2.1 and above. Counts were first performed using an automatic built-in spot detection algorithm in the Imaris software within a pre-set region of interest (ROI). Positive signals within the ROI with a diameter larger than 10 *µ*m that co-localised with DAPI were identified as spots or positive counts. To determine the co-localisation of double-labelled cells, a spot co-localisation algorithm in the Imaris software was used. Two spots were considered co-localised if the distance between their centre of was less than or equal to 6 *µ*m. Potential false positives were then manually validated by visual inspection of the overlapping signals.

### Statistical analysis

Linear regression was used to determine the correlation of two genes in expression datasets, using the R2: microarray analysis and visualization platform (http://r2.amc.nl) as previously described (Bunt et al., 2010). The differential expression of *NFI* between tumour types was compared with a Welch’s t-test on normalised data. To compare the co-staining distribution of NFIA or NFIB with other cell markers between GBM tissues or to determine gene enrichment, a paired one-way ANOVA with Bonferroni’s multiple comparison correction was performed. To compare the co-staining of GFP-positive electroporated cells with markers in GBM xenografts between different plasmids, a Welch’s t-test was performed. These tests were computed with GraphPad Prism 7 (GraphPad Software, San Diego, California).

## Acknowledgements

We thank the staff of the University of Queensland Biological Resources (UQBR) animal facility and the QBI Advanced Microscopy and Analysis Facility for their expertise and assistance in this project. We thank Rowen Tweedale for critical comments on the manuscript and Alan Ho expert assistance with statistical analyses. We thank Andrew W. Boyd and Richard M. Gronostajski for their advice on this project. The primary human GBM samples and de-identified data used in this project were sourced from the Wesley Medical Research Tissue Bank with appropriate ethics approval.

This work was supported by a National Health and Medical Research Council (NHMRC) project grant (GNT1100443 to LJR), Tour de Cure Young Research Grant (to JB), Brain Foundation research gift (to JB) and Ride for Rhonda research gift (to LJR and JB to support CRB). LJR was supported by an NHMRC Principal Research Fellowship (GNT1120615). HA was supported by a University of Malaya research grant (RP049-17HTM). KSC was supported by a University of Queensland (UQ) International Postgraduate Student Scholarship. JWCL was supported by an Australian Government Research Training Program Scholarship and UQ Centennial Scholarship. ZL was supported by a UQ Graduate School Scholarship.

