## Supplementary Figures for "Transcription factors NFIA and NFIB induce cellular differentiation in high-grade astrocytoma"

Supplementary Figure 1

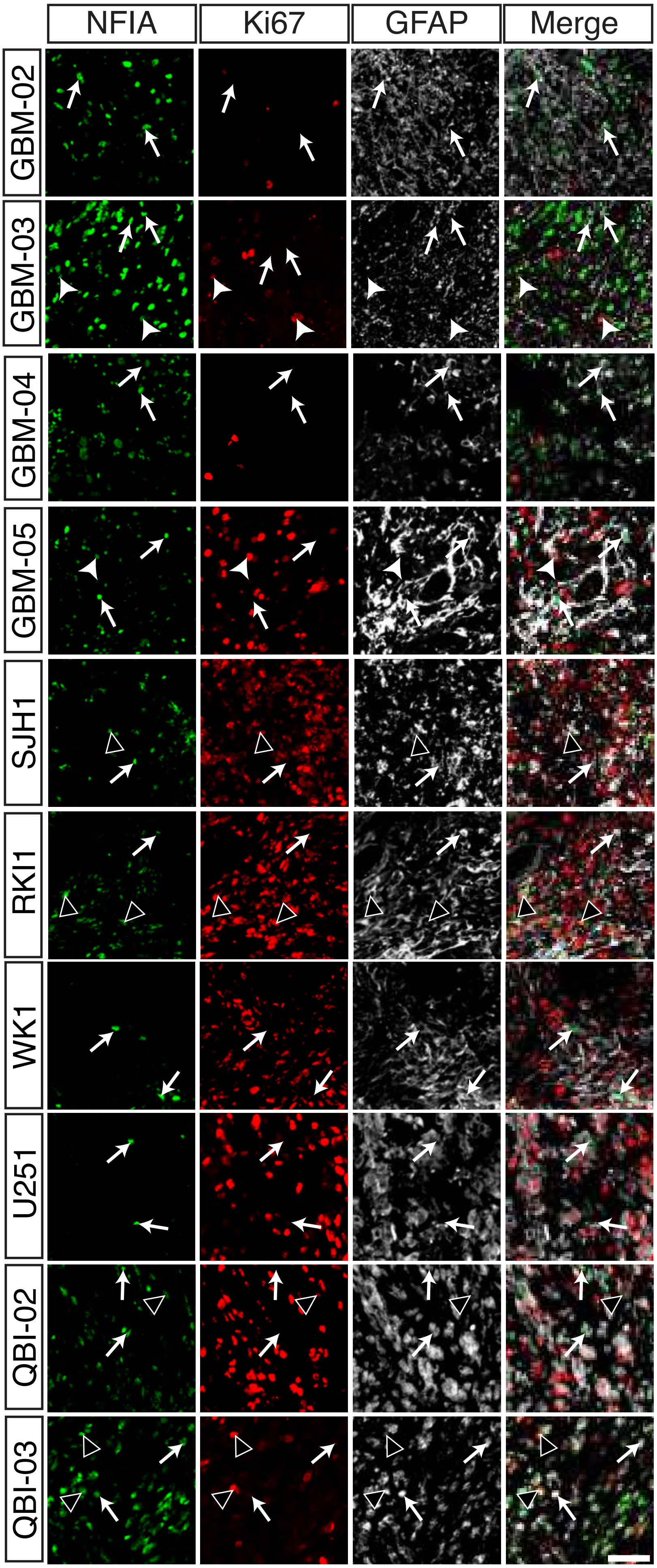

**Supplementary Figure 1** NFI-expressing cells in GBM are predominantly associated with the astrocytic differentiation marker GFAP. Representative images of co-staining of NFIA (green) with GFAP (white) and/or the proliferation marker Ki67 (red) in 11 out of 14 GBM tissues (Fig. 3a). Closed arrowhead: NFI and Ki67 co-localization; open arrowhead: NFI, GFAP and Ki67 co-localization; arrow: NFI and GFAP co-localization. Scale bar = 50 μm.

Supplementary Figure 2

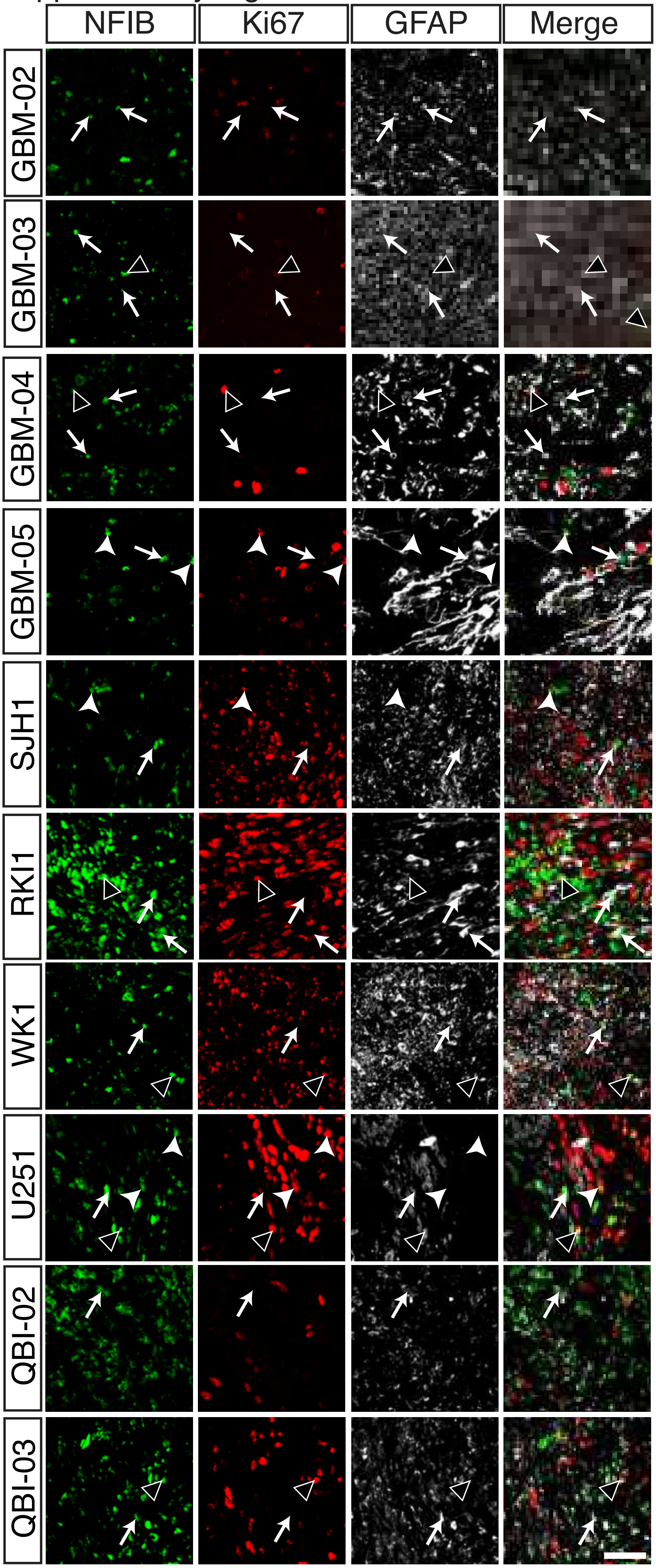

**Supplementary Figure 2** NFI-expressing cells in GBM are predominantly associated with the astrocytic differentiation marker GFAP. **a** and **b** Representative images of co-staining of NFIB (green), with GFAP (white) and/or the proliferation marker Ki67 (red) in 11 out of 14 GBM tissues (Fig. 3b). Closed arrowhead: NFI, GFAP and Ki67 co-localisation; open arrowhead: NFI and Ki67 co-localisation; arrow: NFI and GFAP co-localization. Scale bar = 50 μm.

Supplementary Figure 3

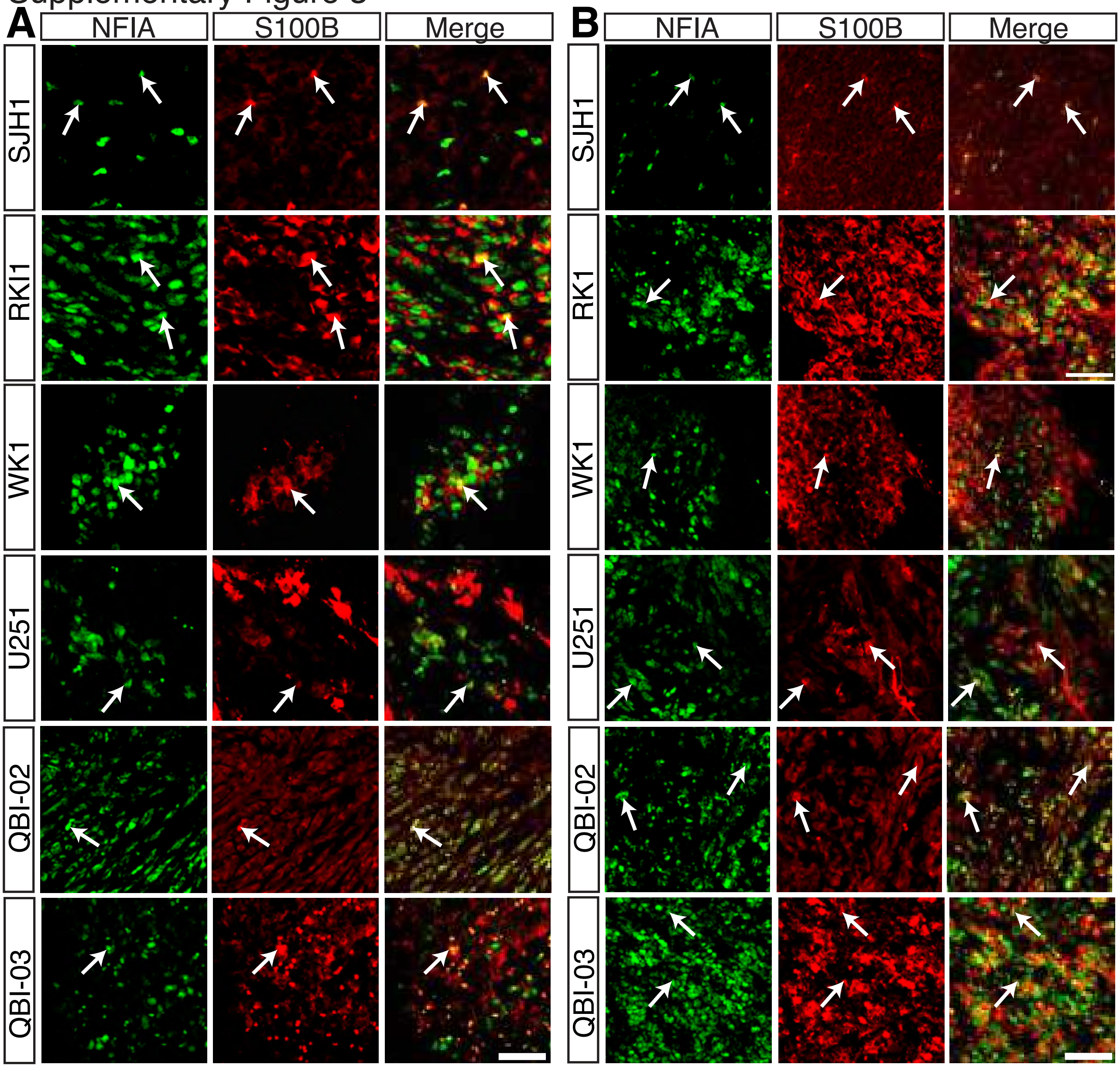

**Supplementary Figure 3** NFI-expressing cells in GBM are predominantly associated with the astrocytic differentiation marker S100B. **a** and **b** Co-staining of NFIA (a; green) or NFIB (b; green), with S100B (red) in six out of eight GBM tissues (Fig. 3c and d). Arrow: NFI, S100B and Ki67 co-localization. Scale bar = 50 µm.

Supplementary Figure 4

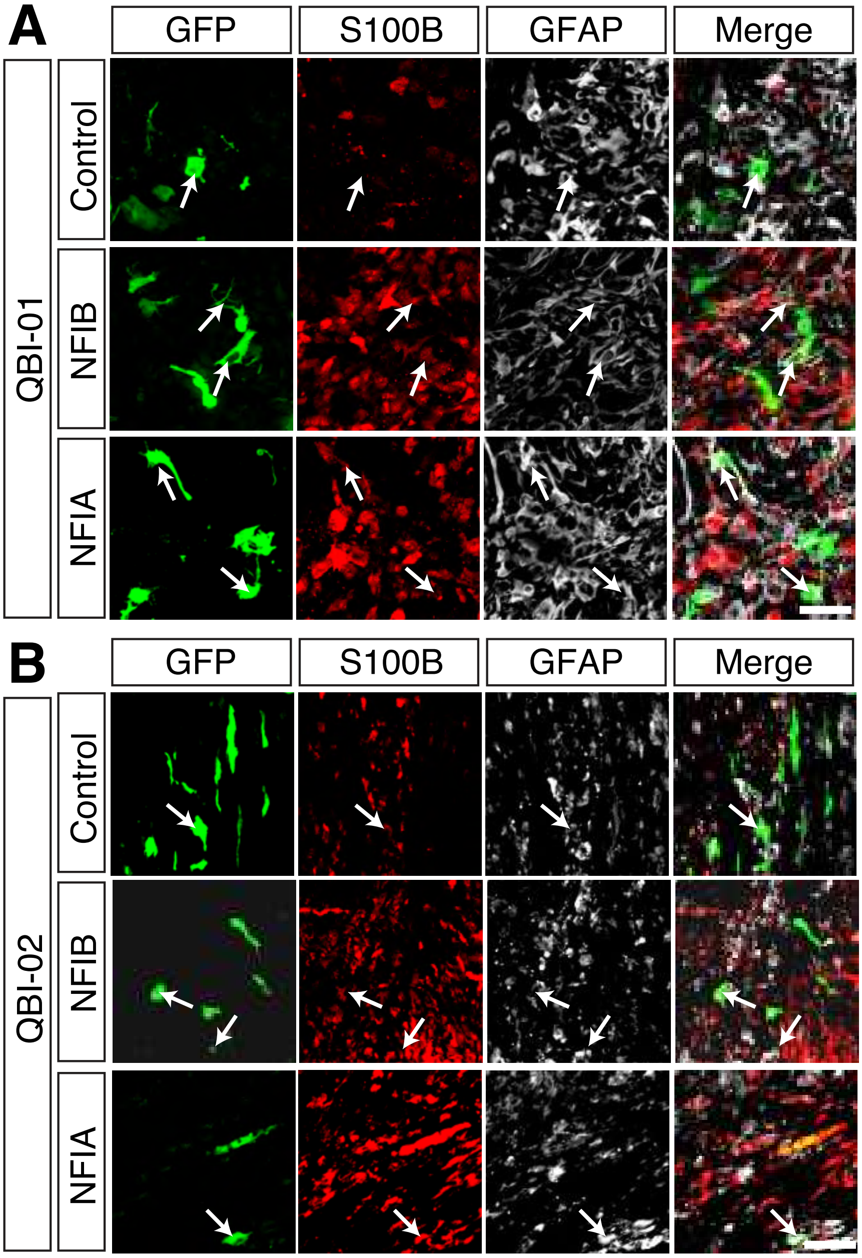

**Supplementary Figure 4** *In vivo* electroporation of GBM xenografts with NFI overexpression constructs co-stained with astrocytic differentiation markers. **a** and **b** Examples of GFP-positive cells (green, arrows) as a marker of cells electroporated with control, NFIA or NFIB overexpression plasmids in QBI-01 (a) and QBI-02 (b) xenografts, stained for S100B (red) and GFAP (white). Scale bar = 50 μm.
